# A Confounder Controlled Machine Learning Approach: Group Analysis and Classification of Schizophrenia and Alzheimer’s Disease using Resting-State Functional Network Connectivity

**DOI:** 10.1101/2023.10.05.561035

**Authors:** Reihaneh Hassanzadeh, Anees Abrol, Godfrey Pearlson, Jessica A. Turner, Vince D. Calhoun

## Abstract

Resting-state functional magnetic resonance imaging (rs-fMRI) has increasingly been used to study both Alzheimer’s disease (AD) and schizophrenia (SZ). While most rs-fMRI studies being conducted in AD and SZ compare patients to healthy controls, it is also of interest to directly compare AD and SZ patients with each other to identify potential biomarkers shared between the disorders. However, comparing patient groups collected in different studies can be challenging due to potential confounds, such as differences in the patient’s age, scan protocols, etc. In this study, we compared and contrasted resting-state functional network connectivity (rs-FNC) of 162 patients with AD and late mild cognitive impairment (LMCI), 181 schizophrenia patients, and 315 cognitively normal (CN) subjects. We used confounder-controlled rs-FNC and applied machine learning algorithms (including support vector machine, logistic regression, random forest, and k-nearest neighbor) and deep learning models (i.e., fully-connected neural networks) to classify subjects in binary and three-class categories according to their diagnosis labels (e.g., AD, SZ, and CN). Our statistical analysis revealed that FNC between the following network pairs is stronger in AD compared to SZ: subcortical-cerebellum, subcortical-cognitive control, cognitive control-cerebellum, and visual-sensory motor networks. On the other hand, FNC is stronger in SZ than AD for the following network pairs: subcortical-visual, subcortical-auditory, subcortical-sensory motor, cerebellum-visual, sensory motor-cognitive control, and within the cerebellum networks. Furthermore, we observed that while AD and SZ disorders each have unique FNC abnormalities, they also share some common functional abnormalities that can be due to similar neurobiological mechanisms or genetic factors contributing to these disorders’ development. Moreover, we achieved an accuracy of 85% in classifying subjects into AD and SZ where default mode, visual, and subcortical networks contributed the most to the classification and accuracy of 68% in classifying subjects into AD, SZ, and CN with the subcortical domain appearing as the most contributing features to the three-way classification. Finally, our findings indicated that for all classification tasks, except AD vs. SZ, males are more predictable than females.

## Introduction

Alzheimer’s disease (AD) represents a neurodegenerative condition and is the most prevalent form of dementia impacting elderly people [1]. Schizophrenia (SZ), on the other hand, is a psychiatric disorder that is characterized by dysconnectivity^1^ between brain regions, commonly develops in adolescence to young adulthood and continues throughout life [2]. Schizophrenia was initially referred to as “dementia praecox” [3], meaning a premature onset of dementia. People who are diagnosed with schizophrenia at a later stage (after the age of 60) tend to have a higher chance of developing dementia in the future [4]. Similarly, [5–8] have reported subjects with schizophrenia are associated with an increased risk of having dementia. In light of this, the researchers in [9] evaluated the white matter deficit patterns and observed similarities in such patterns between AD and SZ, which may explain the increased risk of developing AD in patients with SZ. Furthermore, some psychiatric symptoms are shared between the two disorders [10], and a person with dementia may experience psychotic symptoms despite never having suffered from schizophrenia [11]. All these studies together suggest that schizophrenia and dementia might be connected; however, the nature of this relationship (e.g., similarities and differences in brain patterns) remains unclear, and therefore more research is required.

In recent decades, machine learning (ML) has become increasingly popular as a tool for analyzing neurological and psychiatric brain disorders [12–14]. A machine learning model leverages the unique and shared patterns within and between groups of subjects to predict clinical scores/labels. A major challenge in predicting a clinical score using such methods is the presence of confounding variables (see Fig 1). The presence of confounding factors in data may bias machine learning methods since such methods capture the most noticeable patterns in the data. Thus, it is imperative that confounding factors are removed or controlled before analysis is conducted.

**Fig 1.**
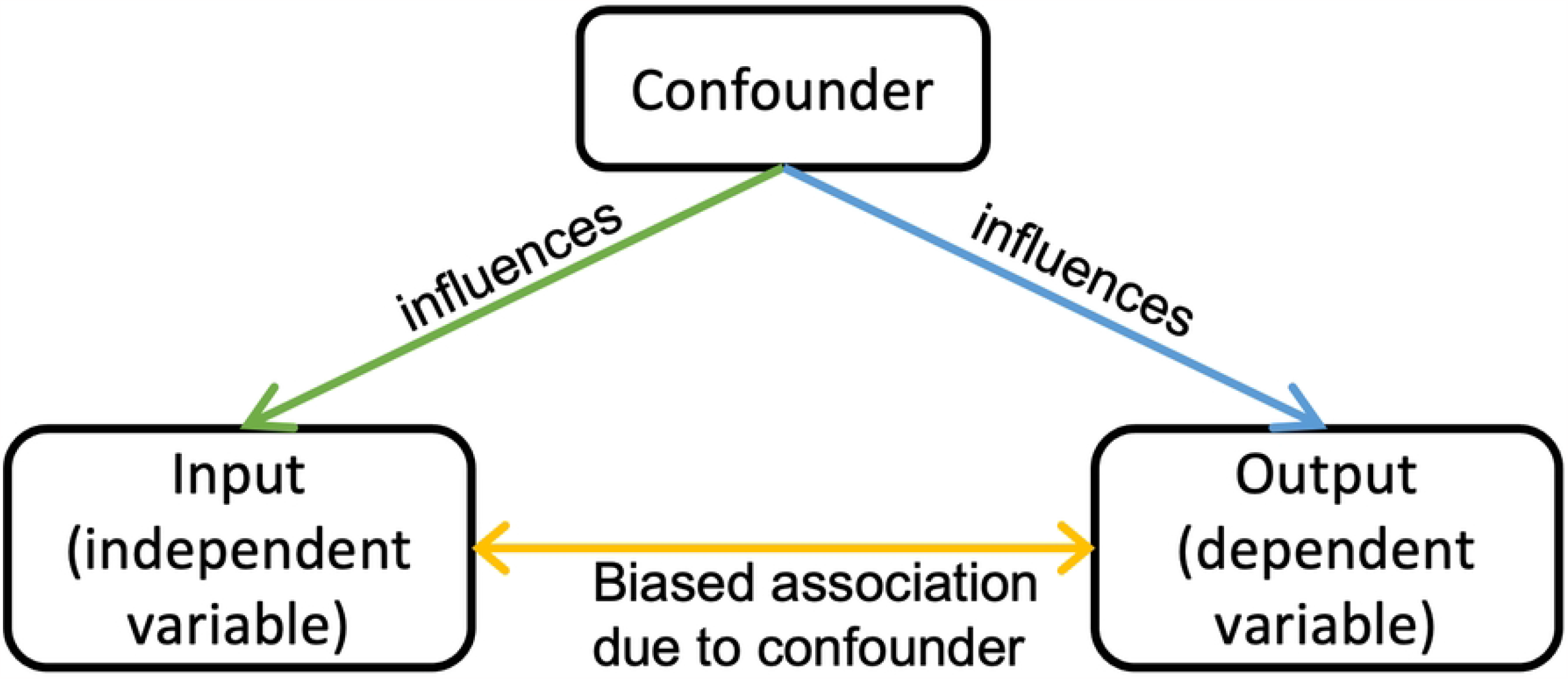
A confounding variable influences the value of both the input and the output and distorts the true relationship between the input and output.

In this paper, we investigated shared and unique functional brain patterns between Alzheimer’s disease and schizophrenia derived from resting-state functional magnetic resonance imaging (rs-fMRI) images [15]. For this purpose, we first measured functional network connectivity (FNC) [16] of the rs-fMRI data using independent component analysis (ICA) [17] and then minimized the effects of three potential main confounders, i.e., age, sex, and site, from the FNC using mathematical and statistical methods. Subsequently, we explored the similar and different patterns in confounder-controlled FNC changes between diagnosis groups (i.e., AD, SZ, and CN subjects) by 1. conducting statistical group comparison experiments and 2. applying machine learning and deep learning (DL) classifiers (including support vector machine, logistic regression, random forest, k-nearest neighbor, and fully-connected neural networks). Lastly, we explored the importance of FNC features and the role of age and sex in classification performance.

## Materials and methods

### Participants

We used rs-fMRI data from the Bipolar and Schizophrenia Network on Intermediate Phenotypes (B-SNIP) [18] available as of 12/09/2014 and from all phases (1/2/go/3) of the Alzheimer’s Disease Neuroimaging Initiative (ADNI)^2^ dataset, available on the ADNI portal on 13/08/2021. Importantly, we did not have access to information that could identify individual participants either during or after data collection. For ADNI, we pooled the AD and late mild cognitive impairment (LMCI) subjects. To balance the ADNI data by diagnosis, we subsampled the control subjects so that the number of control subjects is not more than the number of AD+LMCI subjects within each site. For B-SNIP, we used SZ subjects and controls of the dataset. The final data used in this study consists of 181 SZ subjects and 166 controls from the B-SNIP and 164 AD+LMCI (including 95 AD and 69 LMCI) subjects and 149 controls from ADNI. Table 1 and <u>S1 Fig</u> include demographic information and distribution of the participants used in this study.

**Table 1.** Participants’ demographic characteristics.

|  | Sample size | Age (years): *Mean ± SD* [Min,Max] | Gender: Male |
| --- | --- | --- | --- |
| **ADNI** | 313 (47.6% CN) | 77.2 ± 8 [55.3, 95.8] | 52.4% |
| **B-SNIP** | 345 (48.1% CN) | 36.4 ± 12.5 [15, 64] | 56.6% |

### Image acquisition parameters and preprocessing

All subjects were scanned (eyes-open) by 3.0 Tesla scanners with different models, including Siemens TrioTim, GE Signa HDX, Philips, and Siemens Allegra. ADNI data were collected from 46 locations/sites, and B-SNIP data were collected from 6 sites. The imaging parameters of the two datasets are listed in Table 2.

**Table 2.** Imaging parameters.

|  | Sites | Scanner Vendor/Model | Voxel Size (mm) | Acquisition Matrix (mm) | No of Slices | Flip Angle (°) | Repetition Time (ms) | Echo Time (ms) |
| --- | --- | --- | --- | --- | --- | --- | --- | --- |
| **ADNI** | All sites | Philips | 3.3x3.3x3.3 | 64x64 | 48 | 80 | 1800 | 30 |
| **B-SNIP** | Baltimore | Siemens TrioTim | 3.4x3.4x3 | 64x64 | 36 | 70 | 2210 | 30 |
|  | Boston | GE Signa HDX | 3.4x3.4x5 | 64x64 | 30 | 60 | 3000 | 27 |
|  | Chicago | GE Signa HDX | 3.4x3.4x4 | 64x64 | 29 | 60 | 1775 | 27 |
|  | Dallas | Philips | 3.4x3.4x4 | 64x64 | 29 | 60 | 1500 | 27 |
|  | Detroit | Siemens TrioTim | 3.4x3.4x4 | 64x64 | 29 | 60 | 1570 | 22 |

The fMRI results underwent preprocessing through an SPM12 workflow that encompassed rigid body motion adjustment to rectify participant head movement, synchronization of slice timing, transformation to the standard MNI space utilizing the EPI template, resampling to isotropic voxels of (3mm)3, and smoothing via a Gaussian kernel (FWHM = 6mm). Quality assurance (QA) of the processed fMRI datasets involved eliminating images with low correlation to individual and/or group-level masks. Furthermore, fMRI data containing excessive head motion were removed to prevent possible erroneous variations in functional connectivity.

### Independent component analysis (ICA)

According to the pipeline in Fig 2, the first step of this study was to estimate the brain networks of fMRI images. To do so, we used a spatially-constrained group ICA [17] using the neuromark fMRI 1.0 template [19] (available in the GIFT toolbox http://trendscenter.org/software/gift and also at http://trendscenter.org/data) to estimate 53 spatially independent reproducible brain components and associated time courses (see <u>S1 Table</u> and <u>S2 Fig</u> in Supplementary Information). For further analysis, the 53 brain components were classified into seven functional domains, namely subcortical (SC), auditory (AU), visual (VI), sensorimotor (SM), cognitive control (CC), default mode (DM), and cerebellar (CB). Fig 3 visualizes the spatial maps of the selected ICA components.

**Fig 2.**
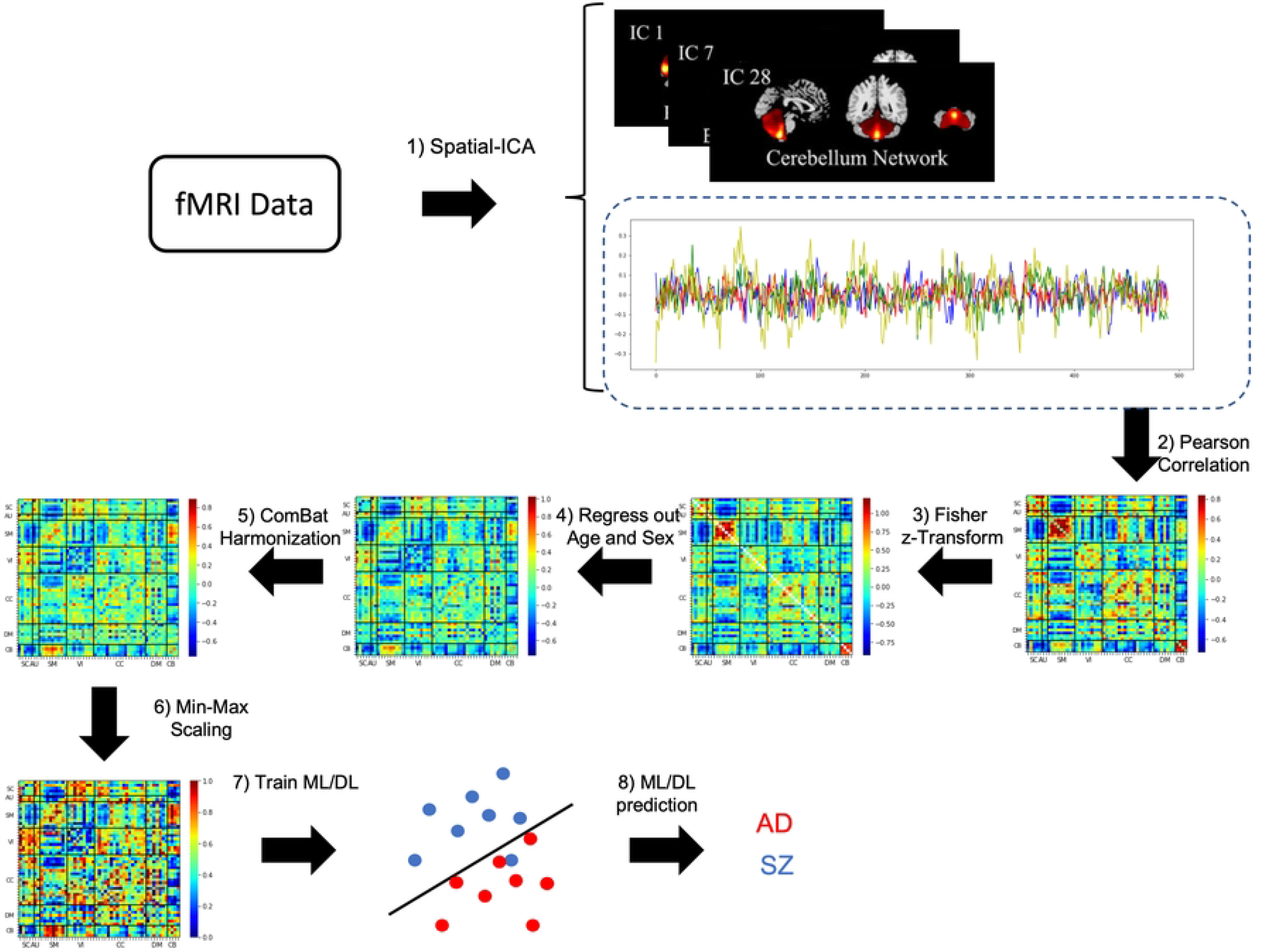
Method pipeline. Initially, we employ a spatially-restricted group ICA to decompose rs-fMRI data into spatial maps and time courses. Following this, we compute FNC features using Pearson’s correlation coefficient for the ICA-derived time courses. Subsequently, we utilize Fisher transformation on the FNC features to stabilize variance, enabling more accurate statistical inferences. We then employ multiple linear regression to eliminate age and gender variability and use an empirical Bayesian approach, specifically ComBat, to eradicate site-related effects from the FNC features. Lastly, we standardize each subject’s FNC matrix via min-max scaling and input the normalized FNC into machine learning classifiers to predict diagnostic outcomes for the subjects.

**Fig 3.**
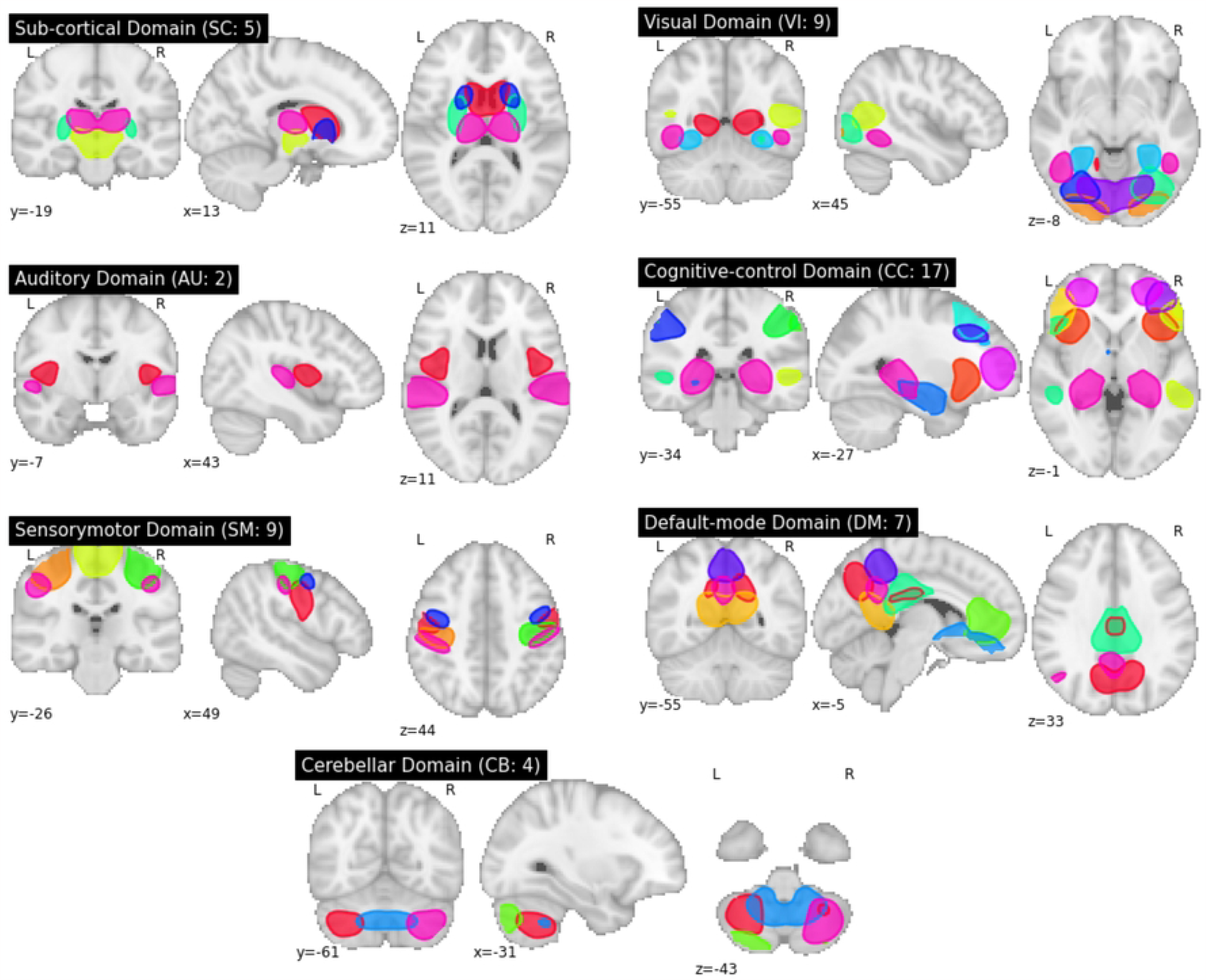
Thresholded spatial maps. 53 functional brain networks are grouped into 5 SC, 2 AU, 7 VI, 7 SM, 14 CC, 7 DM, and 5 CB networks.

### Functional network connectivity (FNC)

An FNC matrix of size 53x53 is calculated for each subject. Each cell in the matrix is a Pearson correlation coefficient between the two corresponding ICA-estimated time courses. We Fisher-transformed the correlation coefficients to stabilize the variance such that it does not depend on the underlying correlation coefficients [20].

### Minimizing confounding effects

Age, gender, and scanning site are three potential confounding variables in our study datasets (see <u>S1 Fig</u> in Supplementary Information). To eliminate age and gender variability across datasets, we applied multiple linear regression to reduce the linear relationship between an FNC feature (i.e., a dependent variable) and age and gender (i.e., independent variables). After regressing out the age and gender information, we performed the well-known site harmonization technique, Combining Batches (ComBat) [21], to mitigate the site effects (i.e., variability due to acquisition parameters). ComBat is an empirical Bayesian method that removes nonbiological variabilities (i.e., site-related effects) while preserving the association between the input (e.g., FNC features) and the target variable (e.g., diagnosis).

### Machine and deep learning classifiers

Machine learning (ML) and fully connected neural networks (NN) classification models have proven their ability to identify and detect variations in the brain among different subject cohorts. In this study, we used four ML methods, including support vector machine (SVM), logistic regression (LR), random forest (RF), and k-nearest neighbor (KNN), and a fully connected neural network consisting of an input layer, one hidden layer, and output layer, for classifying subjects into diagnostic categories. The hyperparameter details for ML and NN models are summarized in <u>S2 Table</u> to <u>S6 Table</u> in Supplementary Information.

## Experiments and results

### Group differences

Two-sample *t*–tests were performed to compare FNCs for the AD+LMCI (henceforth referred to as AD) vs. SZ, AD vs. CN, and SZ vs. CN groups. The false-discovery rate (FDR [22]) was used to control for the multiple testing problem. Fig 4 demonstrates *t*-values (U) and significant *t*-values after passing the FDR with *α, q* = 0.05 (L). According to Fig 4.A, patients with AD have stronger FNC primarily in the SC-CC, SC-CB, SM-VI, and CC-CB networks, while patients with SZ have stronger FNC mostly in the SC-AU, SC-SM, SC-VI, SC-DM, SM-CC, VI-CB, and intra-CB networks. The AD vs. CN group comparison results in Fig 4.B demonstrate increased FNC (i.e., hyperconnectivity) between SC and CC and within SC networks and decreased FNC (i.e., hypoconnectivity) between SC and VI and within CB networks. Moreover, Fig4.C displays the comparison outcomes of SZ against CN, revealing a significant increase in functional connectivity between CB and SM as well as between CB and VI networks but a significant decrease in VI-SM, CB-CC, and CB-SC networks of SZ in comparison to control subjects. Overall, the findings in Fig4 demonstrate that substantial group differences exist between subject cohorts, and these differences seem to be closely related to the specific functional domain involved.

**Fig 4.**
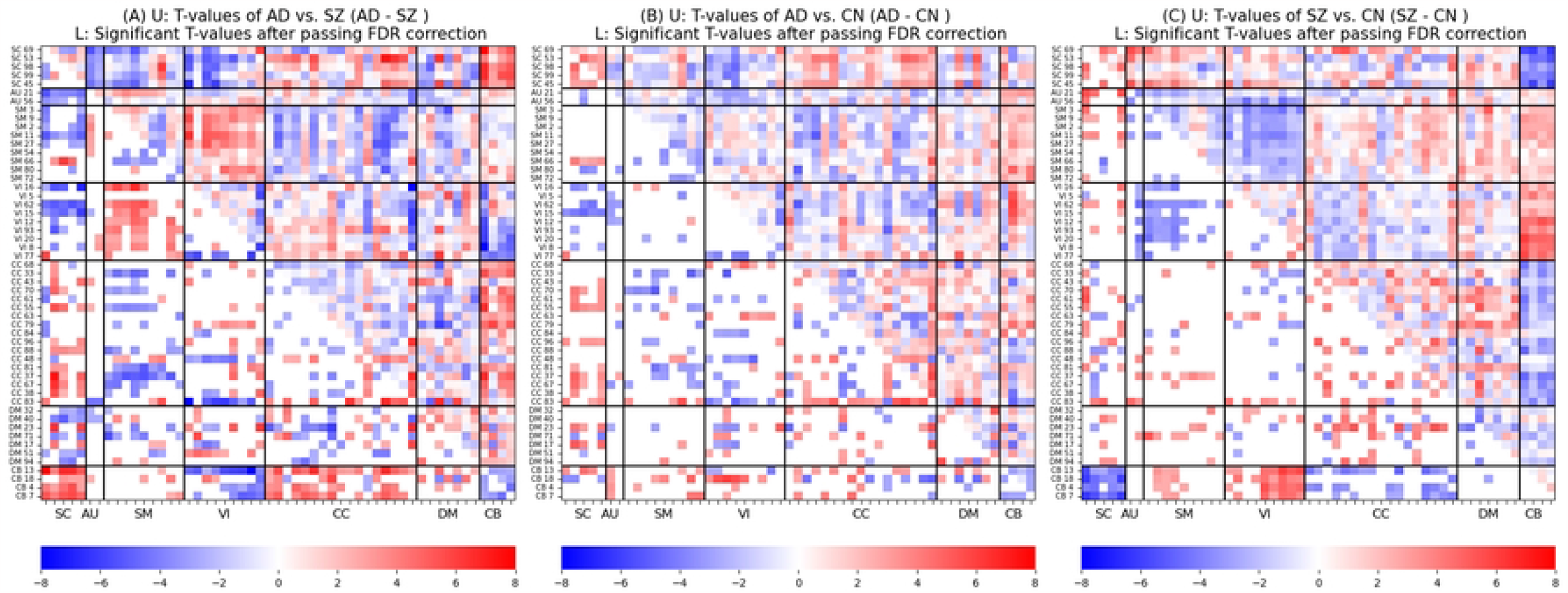
Group differences in FNCs. Upper triangle (U): positive and negative *t*-values from two-sample *t*–tests (*α* = 0.05) are colored in red and blue, respectively. Lower triangle (L): significant *t*-values pass the multiple comparisons, FDR (*q* = 0.05). Fig 4.A suggests that patients with AD have stronger FNC primarily in the SC-CC, SC-CB, SM-VI, and CC-CB networks, while patients with SZ have stronger FNC mostly in the SC-AU, SC-SM, SC-VI, SC-DM, SM-CC, VI-CB, and intra-CB networks. Fig 4.B demonstrates hypoconnectivity between SC and VI and within CB networks and hyperconnectivity between SC and CC and within SC networks. Fig 4.C indicates FNC in SZ increases between CB and VI as well as between CB and SM networks but decreases in VI-SM, CB-CC, and CB-SC networks when compared with controls.

### Common and specific FNC changes between AD and SZ (when compared with CN)

We compared the group differences of AD and SZ against CN (Fig 4.B and Fig 4.C) to each other to identify common and specific FNC abnormalities between the two disorders. According to Fig 5.A-B, common increased FNC was primarily observed in CB 18 (Cerebellum, refer to <u>S1 Tab</u>le), CC 63 (Inferior parietal lobule), CC 83 (Hippocampus), CC 43 (Superior medial frontal gyrus), SC 45 (Thalamus), and SC 98 (Putamen) while common decreased FNC was mainly found in CB 18 and VI 20 (Inferior occipital gyrus).

**Fig 5.**
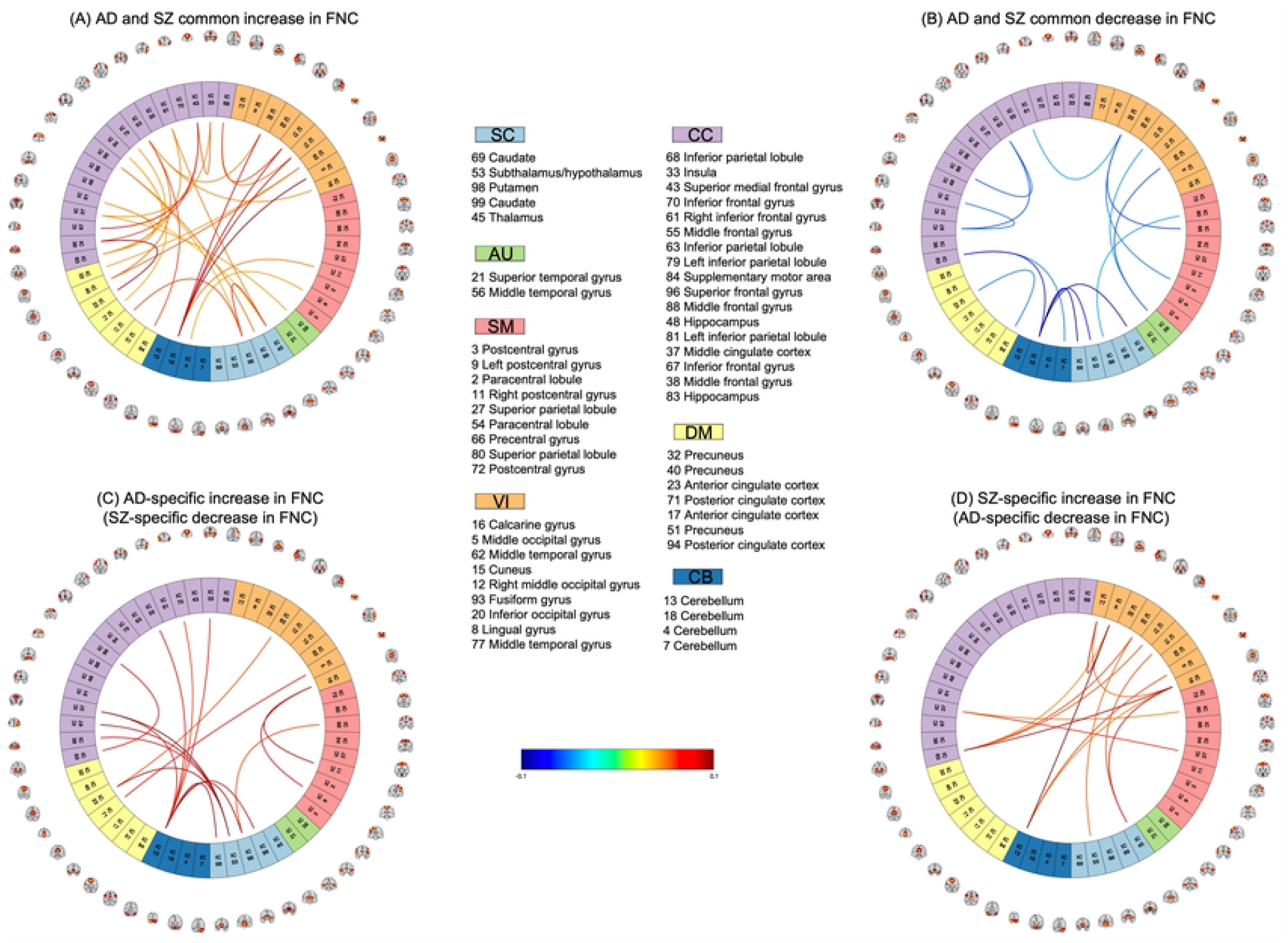
Common and unique FNC changes in AD and SZ when compared with CN. Fig 5.A and Fig 5.B show that common increased FNC is primarily observed in CB 18, CC 63, CC 48, CC 81, CC 68, and SC 98 and decreased FNC is mainly found in CB 18 and VI 20. Figures 4.C and 4.D show the uniqueness mostly occurs in CB 13, SC 53, CC 83, CB 13, VI 16, and VI 8.

Moreover, the AD-unique changes in FNC (Fig 5.C) are noticeably found in CB 13 (Cerebellum) and SC 53 (Subthalamus) and the SZ-unique changes in FNC (Fig 5.D) are primarily observed in CC 83 (Cerebellum), CB 13 (Cerebellum), VI 16 (Calcarine gyrus), and VI 8 (Lingual gyrus). Moreover, we found that the unique changes of both disorders in FNC point to significant group differences in Fig 4.A.

### Classification performance

We implemented a stratified 10-fold cross-validation (CV) strategy where each diagnostic class (i.e., AD, SZ, and CN) is represented (approximately) equally in each set (i.e., training and test sets). We repeated the CV procedure for 3 iterations (30 runs in total) to provide a less noisy estimated performance. At each run, 9 folds were considered for training a model where 10% of the training data was used as a validation set for tuning the hyper-parameters listed in <u>S2 Table</u> to <u>S6 Table</u> in Supplementary Information. Finally, when the optimal model was selected, the remaining one fold was used to test the model. It should be noted that the results reported as part of this section have been averaged over all 30 runs in this study.

Fig 6 shows the accuracy, sensitivity, and specificity of the machine learning algorithms (i.e., SVM, LR, RF, and KNN) and deep learning models (i.e., fully-connected neural networks) for all six possible classification tasks on the ADNI and B-SNIP data - AD vs. SZ vs. CN (i.e., three-way classification), AD vs. SZ (i.e., binary classification of AD and SZ), AD vs. CN, SZ vs. CN, AD vs. CN:B-SNIP, SZ vs. CN:ADNI. The first four classification tasks are the main tasks and the focus of this study, whereas the last two classification tasks are used as the baseline for monitoring the potential impact of biases or otherwise model’s variance with respect to the datasets.

**Fig 6.**
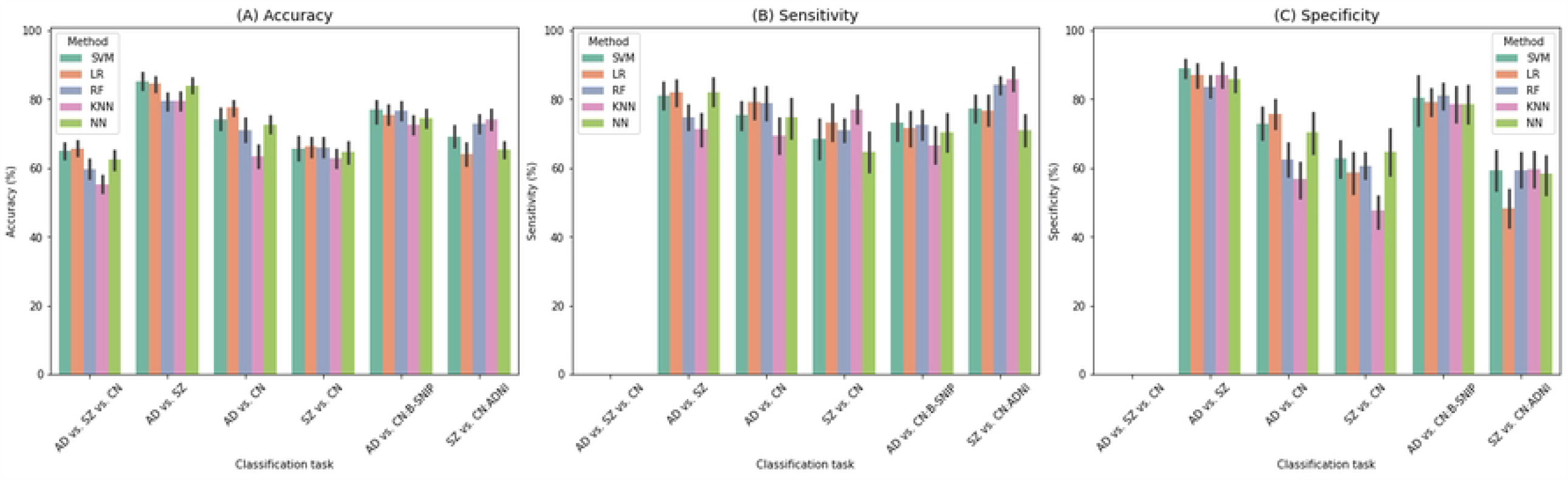
Accuracy, sensitivity, and specificity (*Mean* ± *SE*) in the absence of confounders. According to Fig 6.A, logistic regression (LR) outperforms other models in AD vs. SZ vs. CN, AD vs. CN, and SZ vs. CN; however, SVM performs the best in classifying AD against SZ. Fig 6.B-C show that AD subjects are more predictable than SZ ones, and patients with AD and SZ are more predictable than controls.

According to Fig 6.A, logistic regression resulted in the highest accuracy for the three-way classification of AD, SZ, and CN (68%) and the AD patients against controls (78%) and SZ against controls (66%) classifications, while SVM outperformed the binary classification of AD vs. SZ (85%). In addition, the results demonstrate that machine learning models, especially logistic regression, outperform deep learning models.

Sensitivity and specificity analyses in Fig 6.B and Fig 6.C indicate a higher specificity in the binary classification task (the AD samples were treated as the positive class), suggesting SZ subjects have more unique patterns compared to AD subjects. Moreover, in the AD vs. CN and SZ vs. CN classification tasks, the results show that patients with AD and SZ are more predictable than the control subjects. Lastly, for each classification task, the sensitivity and specificity patterns of each of the machine learning and deep learning models show a similar pattern.

Finally, comparing the baseline results in the absence (Fig 6) and presence <u>(S3 Fig</u> in Supplementary Information) of confounders shows that controlling for confounders is necessary for accurate analysis and conclusions. That is, the amount of variation in accuracy when patients and controls are selected from the same dataset (e.g., AD vs. CN) compared to when they are selected from different datasets (e.g., AD vs. CN:B-SNIP) significantly reduces, which is necessary to have our trained models perform reasonably in a manner that is not dependent on a specific dataset. A large variation between the measured performance for these baselines would suggest that the model is picking up the idiosyncrasies of the datasets rather than capturing the pattern of interest.

### Feature contributions in classification

We used occlusion sensitivity analysis to assess the importance of each brain connection (FNC feature) in the classification of subjects. Specifically, we repeatedly replaced each feature with zero and test the best classifier on the resulting features (i.e., at each run, only one feature is replaced with zero while the values of the remaining features are not changed). Fig 7 shows the top 1% of the most contributing features to the classifiers. According to Fig 7.A, SC 53 (Subthalamus/hypothalamus) is the most contributing network to the three-way classification. Fig 7.B shows that the most important features in the binary classification of AD against SZ subjects are among DM, VI, and SM domains, in particular the DM 23 (Anterior cingulate cortex), VI 77 (Middle temporal gyrus), VI 5 (Middle occipital gyrus), SC 53 (Subthalamus/hypothalamus), and SC 98 (Putamen) networks. Fig 7.C-D show feature importance for classifying patients from controls. These figures highlight that the cognitive-control (e.g., CC 33: Insula) and subcortical (e.g., SC 99: Caudate, SC 98: Putamen) networks play a major role in classifying AD vs. CN and SZ vs. CN, respectively.

**Fig 7.**
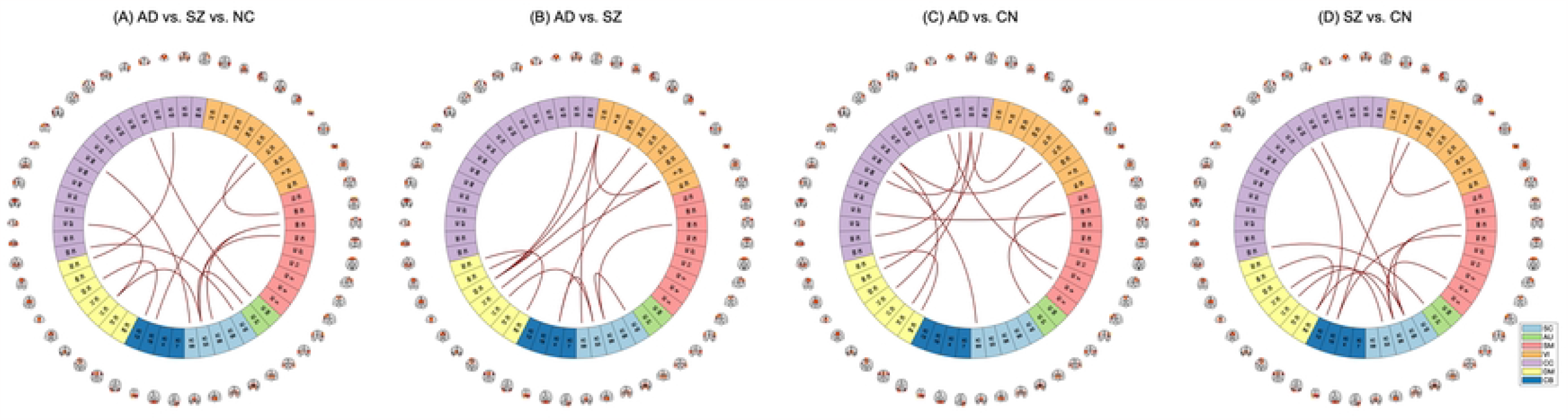
Top 1% important features in classification. (A) Subthalamus/hypothalamus is the most contributing network to the three-way classification. (B) DM, VI, and SM domains contribute the most to the binary classification of AD against SZ subjects, in particular the anterior cingulate cortex, middle temporal gyrus, middle occipital gyrus, subthalamus/hypothalamus, and putamen networks. (C-D) The cognitive-control (e.g., Insula) and subcortical (e.g., Caudate and Putamen) networks play a major role in classifying AD vs. CN and SZ vs. CN, respectively.

### Impact of sex and age on classification accuracy

In order to better understand how demographic information affects classification performance, we divided the test samples based on sex and age and measured the performance of the classifiers for each cohort.

Accordingly, we grouped our subjects based on their gender into females and males. Our results in Fig 8 indicate that males are more predictable than females when classifying patients against controls (i.e., AD vs. CN, SZ vs. CN). Similar founding was observed for the three-way classification (i.e., AD vs. SZ vs. CN), with the male group exhibiting higher accuracy than the female group. Additionally, in the AD vs. SZ classification, it appears that the machine learning model is able to accurately predict the diagnosis of females to a slightly greater extent than that of males.

**Fig 8.**
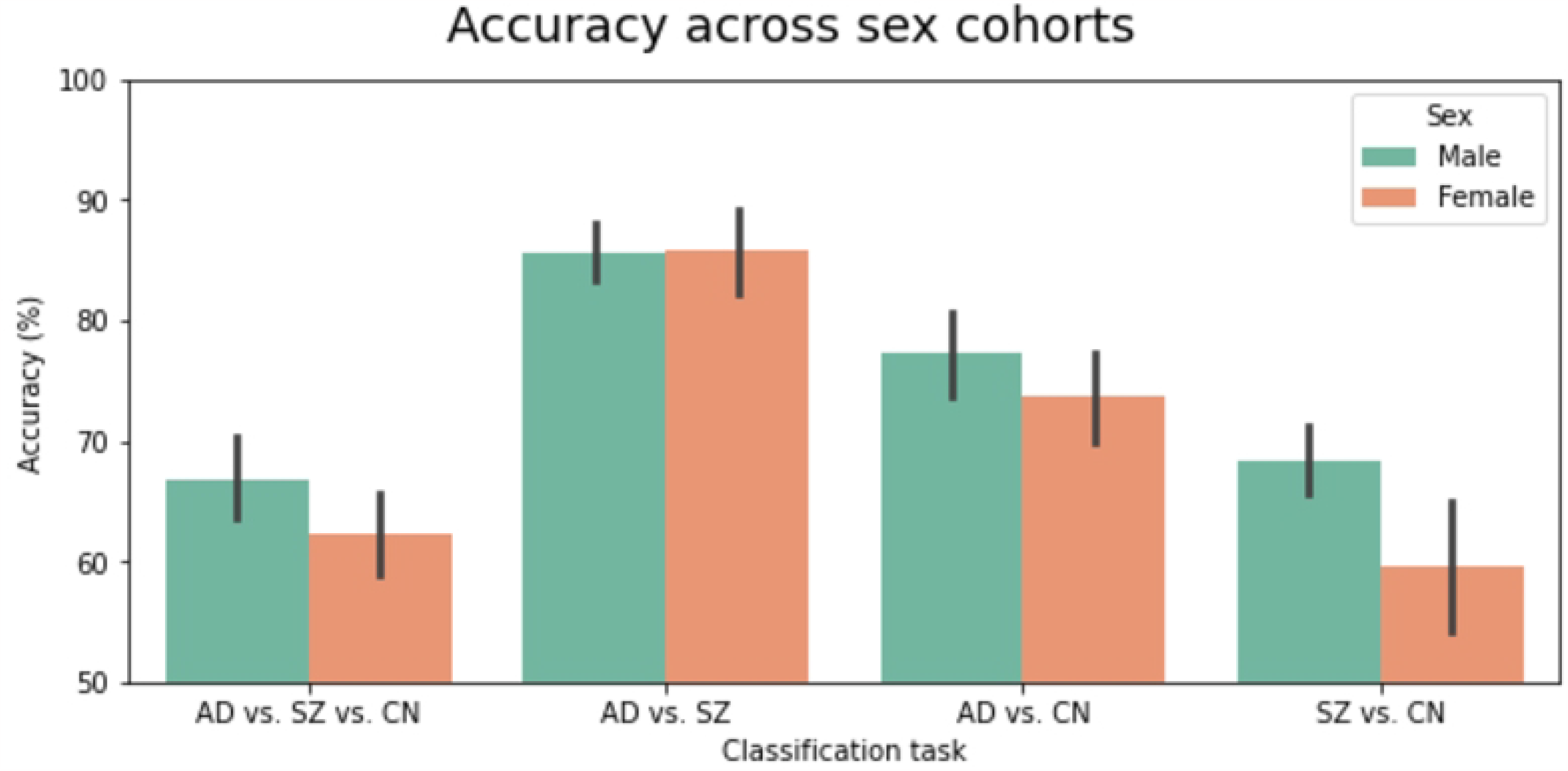
Accuracy (*Mean* ± *SE*) across sex cohorts. According to the figure, males are more predictable than females in the three-way classification and when classifying patients against controls. In the classification of AD vs. SZ, female subjects are more predictable than male subjects.

Additionally, we examined the role that age plays in classification performance. For the three-way and AD vs. SZ classification tasks, each diagnostic group (i.e., AD, SZ, and CN) was divided into young and old cohorts so that young and old cohorts included subjects whose ages were below or above the median of their respective groups.

Fig 9.A-B show that older subjects are more predictable than younger ones for all diagnostic groups (except for controls in the three-way classification where accuracy is approximately the same). In addition, for the disorders against controls classification tasks, the test samples were split based on the median of the dataset, and we observed similar results showing higher accuracy for the older subjects (see Fig 9.C).

**Fig 9.**
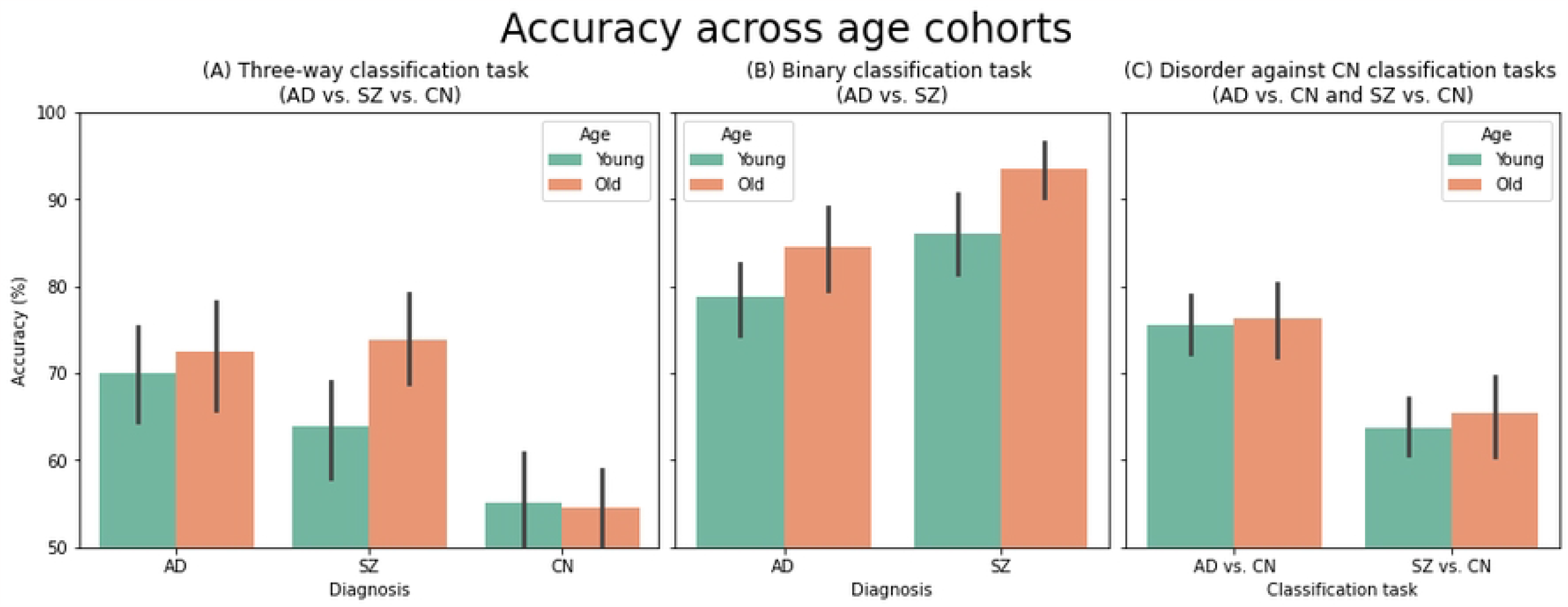
Accuracy (*Mean* ± *SE*) across age cohorts. In all classification tasks, older subjects are more predictable than younger subjects, except for controls in the three-way classification, where accuracy is approximately the same.

## Discussion

Schizophrenia and Alzheimer’s disease are recognized today as distinct brain disorders with different pathologies; schizophrenia is primarily diagnosed based on psychiatric symptoms, while Alzheimer’s is primarily diagnosed based on neurological symptoms. However, there may be some connections between the two disorders. Similar clinical symptoms and brain dysfunction patterns [10] between the two disorders and an increased risk of dementia in schizophrenia [5–8] suggest there may be some functional connectivity overlap between AD and SZ. Therefore, additional research is required to understand how SZ and AD are related and unique in terms of functional abnormalities and patterns in the brain. This research work is, to our knowledge, the first study to investigate the relationship (including its commonality and specificity) between AD and SZ based on functional resting-state FNC.

### Statistical group analysis

Based on the group comparison results between AD and SZ (Fig 5.A), we observed that the functional connectivity between SC and CB, CC and CB, and VI and SM networks are stronger in AD than SZ. Conversely, connectivity between CB and VI (e.g., between calcarine gyrus and hippocampus), AU and SC, and SC and VI (e.g., between calcarine gyrus and thalamus), and within CB networks, are weaker in AD.

In addition, when compared to the previous works, we found consistent patterns in FNC when comparing AD and SZ to controls (Fig 5.B and Fig 5.C). For example, in SZ vs. CN, we observed hypo-connectivity between CB and SC networks (consistent with observations in [19]) and hyper-connectivity within SC networks (consistent with [23]) and between CB and SM networks (consistent with findings reported in [19]). For AD compared to the control group, we found decreased FNC within the left inferior parietal lobule networks (i.e., between IC 79 and IC 81) [24], within the hippocampus networks (i.e., between IC 48 and IC 83) [24, 25], which are one of the first regions in the brain to be affected by AD [26, 27], between the posterior cingulate cortex (i.e., IC 71 and IC 94) and hippocampus networks (i.e., between IC 71 and IC 83) [28], between the temporal lobe (including superior temporal gyrus, middle temporal gyrus, and middle temporal gyrus) and corpus striatum (i.e., caudate, putamen, caudate) [29], and between the thalamus (i.e., subthalamus/hypothalamus and thalamus) and occipital lobe (i.e., visual networks) [29]. Additionally, we observed increased connectivity between the frontal lobe (i.e., the frontal gyrus networks) and the corpus striatum (i.e., caudate, putamen, caudate) consistent with the findings in [29].

### Shared and unique abnormal FNC patterns in AD and SZ

We observed schizophrenia and Alzheimer’s disease had both common and distinct abnormalities in functional connectivity when compared to the control group. Interestingly, AD and SZ showed more common FNC abnormalities (46 connections) than unique ones (31 connections), further supporting the hypothesis that these conditions are related [4–8].

Most of the common abnormalities included an increased change in FNC, which mainly occurred between the cerebellum and visual networks, within cognitive control networks, between Inferior parietal lobule and virtual networks and default mode networks, within the subcortical networks, and between the subcortical and cognitive control networks. Common decreased FNC was mainly found between Cerebellum and subcortical networks, between inferior frontal gyrus and parietal lobule networks, within the middle temporal gyrus network, and between inferior occipital gyrus and sensory-motor networks.

Interestingly, we observed that the cerebellum (i.e., IC 18) shows different patterns in connection with the networks of different functional domains. That is, while FNC between the cerebellum and the subcortical networks (including caudate, subthalamus, and thalamus) shows decreased changes in both AD and SZ, the connectivity between the cerebellum and the visual networks (including middle occipital gyrus, cuneus, right middle occipital gyrus, inferior occipital gyrus) reveals increased changes.

Moreover, the AD-unique changes in FNC are noticeably found between the cerebellum (i.e., IC 13) and cognitive control network (including middle cingulate cortex, middle frontal gyrus, middle frontal gyrus, and left inferior parietal lobule) and subcortical network (including ICs caudate, subthalamus/hypothalamus, putamen, and caudate), between and Subthalamus and cognitive control networks (including middle cingulate cortex, middle frontal gyrus, and inferior frontal gyrus) and anterior cingulate cortex. The SZ-unique changes in FNC are primarily observed between the cerebellum (i.e., IC 83) and visual networks, between the cerebellum (IC 13) and visual network, and within visual networks (i.e., between middle temporal gyrus, ICs lingual gyrus, and calcarine gyrus). Additionally, we found that the unique changes of both disorders in FNC point to significant group differences computed from the *t*–test.

Lastly, comparing the common and unique changes shows that networks from the same domain may show different FNC polarity from networks of another domain. For example, while CB 18 and CB 13 are from the same domain, CB 18 appears as a notable network in common FNC changes, but CB 13 reveals unique FNC changes in the two disorders, AD and SZ.

### ML classification

A high accuracy of 85% was achieved in the classification of AD and SZ, suggesting the functional connectivity patterns are largely unique to the disorders. We observed that the default mode, visual, and subcortical functional domains have the most contributing features in classifying AD vs. SZ subjects (see Fig 7.B). Furthermore, most of the features that were deemed most important to the classification also exhibited substantial group differences, as depicted in Fig 4.A. In the case of the three-way classification of AD, SZ, and control (CN) subjects, our approach achieved an accuracy of 68%. In this scenario, the subcortical domain emerged as the most critical functional domain for the classification.

The higher specificity than the sensitivity of the machine learning models in classifying AD against SZ, where AD samples were treated as the positive class, suggests that SZ information can be captured more effectively than AD data, suggesting more distinct functional patterns in patients with SZ.

Furthermore, our observations indicated that machine learning (ML) methods outperformed neural networks in all classification tasks. This can be attributed to the fact that deep models, including neural networks, generally require a substantial amount of data to be trained effectively. Consequently, in situations where only a small sample size is available, ML models may be better suited for learning from a limited amount of data, since they tend to be more efficient in doing so.

One important component of this study was controlling the effects of confounders. If significant confounding variables exist, machine learning models may instead focus more on capturing the variability driven by these variables. We considered age, gender, and site ID as three main confounders and reduce their effects using regression and harmonization approaches. Our results indicate that we achieve similar accuracy on the confounder-controlled inputs whether ML methods train on subjects and controls of the same or different datasets. In contrast, when confounders are not removed, we observe a 99% accuracy in classifying subjects and controls selected from different datasets, suggesting the classifiers are biased by confounding variables. This observation corroborates the necessity of removing confounders before analyzing and predicting.

### Limitations and future directions

In this study, although we minimized three well-known potential confounders, i.e., age, gender, and multisite effects, the effect of confounding was not entirely removed. This might be because of other confounder variables, such as the race of subjects that are not collected for our datasets. Furthermore, as an interesting future direction, one can apply more advanced approaches (e.g., generative adversarial networks (GANs) as suggested by) to control confounders simultaneously to predict the target values (i.e., diagnostic). Finally, the scope of future work on this topic can extend to using other brain features, such as dynamic FNC, ICA-estimated time courses, and even voxel-level MRI images to identify similarities and differences between AD and SZ.

## Conclusion

The aim of this research study was to identify potential biomarkers that are shared and linked between neurodegenerative (i.e., AD) and psychiatric (i.e., SZ) groups. To achieve this, we utilized confounder-controlled resting-state functional network connectivity (FNC) to compare AD and SZ patients. The study involved a statistical group analysis and the use of machine/deep learning models to analyze the subjects based on their diagnosis labels. The results of the study revealed that although each disorder had unique FNC abnormalities, there were also common functional abnormalities that could be due to similar neurobiological mechanisms or genetic factors contributing to these disorders’ development.

We achieved an accuracy of 85% in classifying subjects into neurodegenerative and psychiatric groups, and 68% in classifying subjects into neurodegenerative, psychiatric, and control individuals. Interestingly, we found that males were more predictable than females in classifying subjects. Overall, this research provided valuable insights into the shared and unique functional brain patterns between neurodegenerative and psychiatric groups. It also highlighted the importance of minimizing confounds when comparing patient groups collected in different studies.

In conclusion, this study’s findings suggest that there are potential biomarkers shared between Alzheimer’s disease and schizophrenia, which can contribute to some similar neurobiological mechanisms or genetic factors contributing to these disorders’ development. This research provides new insight into the complex relationship between these two disorders and may pave the way for future research to identify new therapeutic targets that could benefit both patient populations.

## Supporting information

**S1 Fig. Distribution of age, gender, and site ID across datasets**. (A-B) demonstrate that SZ subjects are considerably younger than AD subjects. (C) reveals that the male and female counts within a dataset are unequal, particularly in B-SNIP. (D) highlights that the number of sites in ADNI is significantly greater than in B-SNIP.

**S2 Fig. ICA-driven spatial maps, (the Neuromark templates)**.

**S3 Fig. Accuracy, sensitivity, and specificity (***Mean* ± *SE***) in the presence of confounders**. Comparing the results of AD vs. CN and SZ vs. CN to AD vs. CN:B-SNIP and SZ vs. CN:ADNI suggest that the ML and NN models are biased by the confounders.

**S1 Table. Networks ID, domain, and region identified by ICA**.

**S2 Table. Hyperparameters of support vector machine (SVM)**.

**S3 Table. Hyperparameters of logistic regression (LR)**.

**S4 Table. Hyperparameters of random forest (RF)**.

**S5 Table. Hyperparameters of k-nearest neighbor (KNN)**.

**S6 Table. Hyperparameters of neural network (NN)**.

## Footnotes

1 abnormal/aberrant connectivity

2 http://adni.loni.usc.edu

